# The *fnr-like* mutants enhance isoxaben tolerance by initiating mitochondrial retrograde signalling

**DOI:** 10.1101/2024.01.11.575284

**Authors:** Ronan Broad, Michael Ogden, Peter M Dracatos, James Whelan, Staffan Persson, Ghazanfar Abbas Khan

**Affiliations:** Department of Animal, Plant and Soil Sciences, School of Agriculture, Biomedicine and Environment, La Trobe University, Bundoora, VIC 3086, Australia; Copenhagen Plant Science Center, Department of Plant & Environmental Sciences, University of Copenhagen, Frederiksberg C 1871, Denmark; College of Life Science, Zhejiang University, Hangzhou 310058, China; Joint International Research Laboratory of Metabolic and Developmental Sciences, State Key Laboratory of Hybrid Rice, School of Life Sciences and Biotechnology, Shanghai Jiao Tong University, Shanghai 20040, China

## Abstract

Isoxaben is a pre-emergent herbicide used to control broadleaf weeds. While the phytotoxic mechanism is not completely understood, isoxaben interferes with cellulose synthesis. Certain mutations in cellulose synthase complex proteins can confer isoxaben tolerance; however, these mutations cause compromised cellulose synthesis and perturbed plant growth, rendering them unsuitable as herbicide tolerance traits. We conducted a genetic screen to identify new genes associated with isoxaben tolerance by screening a selection of *Arabidopsis thaliana* T-DNA mutants. We found that mutations in a *FERREDOXIN-NADP(+) OXIDOREDUCTASE-LIKE (FNRL)* gene enhanced tolerance to isoxaben, exhibited as a reduction in primary root stunting, reactive oxygen species accumulation, and ectopic lignification. The *fnrl* mutant did not exhibit a reduction in cellulose levels following exposure to isoxaben, indicating that FNRL operates upstream of isoxaben-induced cellulose inhibition. In line with these results, transcriptomic analysis revealed a highly reduced response to isoxaben treatment in *fnrl* mutant roots. The *fnrl* mutants displayed constitutively induced mitochondrial retrograde signalling, and the observed isoxaben tolerance is partially dependent on the transcription factor ANAC017, a key regulator of mitochondrial retrograde signalling. Moreover, FNRL is highly conserved across all plant lineages, implying conservation of its function. Notably, *fnrl* mutants did not show a growth penalty in shoots, making FNRL a promising target for biotechnological applications in breeding isoxaben tolerance in crops.

## Introduction

Isoxaben (N-[3(1-ethyl-1-methylpropyl)-5-isoxazolyl]) is primarily used as a pre-emergent herbicide for controlling broadleaf weeds (Huggenberger and Gueguen, 1987). Remarkably potent, isoxaben typically displays IC50 values in the nanomolar range (Heim et al., 1989). While the phytotoxic mechanism of isoxaben is not fully understood (Ogden et al., 2023), it significantly reduces the incorporation of radio-labelled glucose into crystalline cellulose (Heim et al., 1990). Cellulose is the main load-bearing component of plant cell walls, consisting of a long chain of β-1,4-linked D-glucose units (Polko and Kieber, 2019; Wang et al., 2020). Cellulose undergoes complex cross-linking with other soluble matrix polysaccharides, such as hemicelluloses and pectin. This network of polymers collectively renders cell walls semipermeable and dynamically adaptable structures (Cosgrove, 2022). Cellulose is synthesised at the plasma membrane by cellulose synthases (CESAs) organised into cellulose synthase complexes (CSCs) (Paredez et al., 2006; Atanassov et al., 2009). The CSC has a distinct structure resembling a rosette with six lobes (Haigler and Brown, 1986), with each lobe predicted to contain three catalytic CESA subunits (Persson et al., 2007).

In Arabidopsis, 10 CESA paralogues are involved in cellulose synthesis, categorised into primary (CESAs 1, 2, 3, 5, 6, and 9, as well as possibly 10) and secondary (CESAs 4, 7, and 8) cell wall cellulose synthesis (Persson et al., 2007; Gonneau et al., 2014). Impaired CESA function leads to a range of growth defects in plants, including loss of anisotropic growth, ectopic lignification, and stunted growth (Arioli et al., 1998; Scheible et al., 2001). Knockout mutants of CESA1 or CESA3 display gametophytic lethality (Persson et al., 2007); however, knockout mutants for other primary cell wall CESAs do not cause lethal phenotypes unless multiple CESAs are dysfunctional, suggesting functional redundancy (Desprez et al., 2007; Persson et al., 2007).

Stress responses to cell wall damage (CWD), induced by the cellulose biosynthesis inhibitor (CBI) isoxaben, have been extensively investigated (Denness et al., 2011; Engelsdorf et al., 2018; Gigli-Bisceglia et al., 2018). These responses are influenced by osmotic support and encompass upregulation of stress response genes, generation of reactive oxygen species (ROS), accumulation of phytohormones like jasmonic acid (JA) and salicylic acid (SA), alterations in cell wall composition, such as ectopic lignification and callose deposition, and eventual growth arrest (Denness et al., 2011; Engelsdorf et al., 2018; Gigli-Bisceglia et al., 2018). Comparable effects have been observed in various *cesa* mutants, providing further insight into the phenotypic manifestations of CWD (Arioli et al., 1998; Fagard et al., 2000).

Plants have developed intricate mechanisms to respond to CWD. Collectively known as cell wall integrity (CWI) pathways, they allow plants to detect and effectively respond to cell wall disruption, such as during cellulose biosynthesis inhibition (Vaahtera et al., 2019). The CWI response to CBIs involves the participation of at least three receptor-like kinases (RLKs) (Vaahtera et al., 2019). One member of the CrRLK1L family, THESEUS1 (THE1), was initially linked to this process as it restored growth of *cesa6* mutants without recovering the cellulose content (Hématy et al., 2007). Furthermore, THE1 is crucial for the altered expression of several stress-response genes when liquid-grown seedlings were exposed to isoxaben (Denness et al., 2011). More recently, two receptor kinases, MALE DISCOVERER1-INTERACTING RECEPTOR LIKE KINASE 2/LEUCINE-RICH REPEAT KINASE FAMILY PROTEIN INDUCED BY SALT STRESS (MIK2/LRR-KISS) (Van der Does et al., 2017) and FEI1/2 (Xu et al., 2008), were implicated in the cell wall stress response induced by isoxaben. Genetic analysis demonstrated that THE1 and MIK2/LRR-KISS share some functions while also having distinct roles, indicating complex regulation of the CBI response (Van der Does et al., 2017).

Apart from receptor-like kinases, mitochondrial retrograde signalling has emerged as a recent player in the CBI response (Hu et al., 2016). Stress conditions typically induce the accumulation of ROS, leading to oxidative damage in mitochondria (Rhoads et al., 2006). Moreover, mitochondria are an important source of cellular ROS (Rhoads et al., 2006; Noctor and Foyer, 2016; Mittler et al., 2022). Stress-induced inhibition of mitochondrial electron transport chain complexes can lead to the non-enzymatic single-electron reduction of oxygen, resulting in the generation of superoxide (O2^•−^), a potent ROS, that is then rapidly converted to hydrogen peroxide (H2O2) by superoxide dismutase and represents the chief messenger molecule in ROS signalling from mitochondria (Noctor and Foyer, 2016; Mittler et al., 2022). As H2O2 is also produced as a signalling molecule in response to a variety of external perturbations, it links external and internal cues to integrate signalling pathways to optimise growth to prevailing conditions (Wu et al., 2020). Perturbations in mitochondrial functions activate signalling cascades from the mitochondria to the nucleus, affecting global gene expression through the mitochondria-nucleus retrograde signalling pathway (Ng et al., 2013; Ng et al., 2014). In plants, several signalling components involved in this process have been identified, including cyclin-dependent kinase E1 (CDKE1) (Ng et al., 2013), as well as the transcription factors ABSCISIC ACID INSENSITIVE4 (Giraud et al., 2009), WRKY40 (Van Aken et al., 2013), NO APICAL MERISTEM/ARABIDOPSIS TRANSCRIPTION FACTOR/CUP-SHAPED COTYLEDON013 (ANAC013) (De Clercq et al., 2013), and ANAC017 (Ng et al., 2013). Depletion of either ANAC013 or ANAC017 disrupts mitochondrial retrograde signalling, resulting in heightened sensitivity to abiotic stresses (De Clercq et al., 2013; Ng et al., 2013), emphasising the significance of mitochondria in stress adaptation.

To identify new components associated with isoxaben resistance, we performed a targeted genetic screen of Arabidopsis T-DNA mutants and found that mutations in a plastid localised *FERREDOXIN-NADP(+) OXIDOREDUCTASE-LIKE* (*FNRL*, At1g15140) conferred isoxaben tolerance. The mutations failed to restore the cellulose deficiency response observed in the *cesa6*/*procuste1* (*prc1-1*) mutant, indicating that *fnrl* is not involved in the CWD response caused by cellulose synthesis inhibition. Moreover, *fnrl* mutants did not show decreased cellulose synthesis in response to isoxaben exposure, suggesting that FNRL functions upstream of the isoxaben-mediated decrease in cellulose synthesis. Furthermore, our investigation revealed that *fnrl* mutants show an induced mitochondrial retrograde signalling response that is partially responsible for the exhibited isoxaben tolerance. Notably, *fnrl* mutants did not exhibit a growth penalty in shoots, making FNRL a promising candidate for biotechnological applications in breeding isoxaben tolerance in crop plants.

## Results

### *fnrl* mutants confer isoxaben resistance and regulate root architecture

To identify genes associated with isoxaben tolerance, we screened homozygous T-DNA *Arabidopsis thaliana* mutants for isoxaben tolerance. Mutants were grown vertically on 1/2 Murashige and Skoog (MS) media supplemented with 2.5 nM isoxaben. A T-DNA line affecting FNRL (SALK_039715, *fnrl-1*) conferred strong isoxaben tolerance as compared to the wild type (Col-0) (Fig. 1A,B; Supplemental Figure 1). The tolerance phenotype was further validated using a second mutant allele of *fnrl* (SALK_048627, *fnrl-2*) and *fnrl-1* complementation with FNRL:GFP expression driven by the endogenous *FNRL* promoter, confirming that *FNRL* is the causal gene responsible for the increased isoxaben tolerance (Fig. 1A,B; Supplemental Figure 1A). The *fnrl-1* allele carries a T-DNA insertion in the first exon, while *fnrl-2* has an insertion in the first intron. Both alleles show no full-length *FNRL* transcript, confirming that both insertions result in gene knockout (Supplemental Figure 1B,C). Furthermore, *fnrl* mutants did not exhibit a reduction in cellulose content upon exposure to isoxaben, further corroborating their tolerance to the isoxaben-mediated inhibition of cellulose synthesis (Fig. 1C).

**Figure 1:**
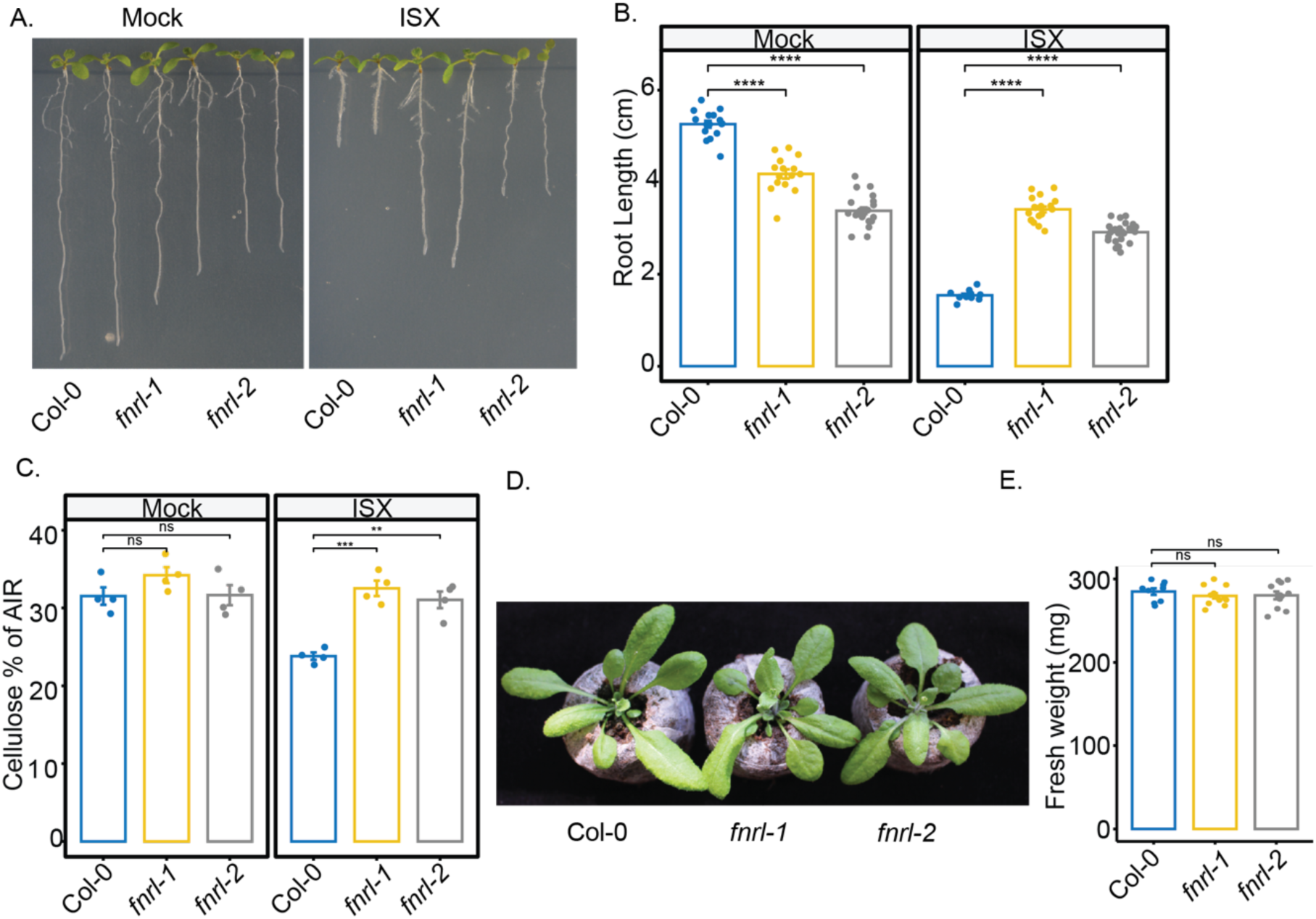
*fnrl* mutants exhibit isoxaben tolerance and modulated root architecture. A) Representative images of ten-day-old seedlings grown on 1/2 MS medium supplemented with 1% sucrose, 0.8% agar, and either 2.5 nM isoxaben (ISX) or Mock treatment. B) Quantification of primary root length in ten-day-old seedlings grown under 2.5 nM isoxaben (ISX) or Mock conditions. C) Cellulose quantification in seedling roots under 2.5 nM isoxaben (ISX) or Mock conditions. AIR: alcohol insoluble residue. D) Representative images of four-week-old rosettes from wild-type and *fnrl* mutant plants grown in soil. E) Rosette weight of four-week-old plants grown in soil. Asterisks denote statistical significance (*, P < 0.05; **, P < 0.01; ***, P < 0.001; and ****, P < 0.0001) according to Student’s t test. The graphics and statistical analysis were generated using the ggpubR package in R.

Interestingly, we observed that both *fnrl* alleles displayed altered root architecture under mock conditions. Specifically, the mutant seedlings exhibited reduced primary root length compared to wild-type plants (Fig. 1A,B). By contrast, rosette growth and fresh weight of the *fnrl* mutants was similar to that of the wild type, suggesting that FNRL mainly affects root architecture rather than overall plant growth and nutrient acquisition (Fig. 1D,E).

### FNRL plays a crucial role in isoxaben-induced cell wall damage response

Typically, isoxaben treatment leads to the accumulation of ROS and lignin in roots (Denness et al., 2011); therefore, we examined these processes in *fnrl* mutants relative to WT in response to isoxaben treatment. We cultivated plants on 1/2 MS plates supplemented with 2.5 nM isoxaben or mock for ten days. To quantify ROS levels, we utilised the cell-permeant indicator 2’,7’-dichlorodihydrofluorescein diacetate (H2DCFDA). The *fnrl* mutants showed reduced ROS presence in the roots compared to the wild type under normal conditions. When wild-type plants were treated with isoxaben, there was a significant increase in H2DCFDA signal intensity compared to the mock treatment, indicating higher ROS accumulation (Fig. 2A,B). However, this isoxaben-induced elevation of H2DCFDA signal was noticeably reduced in both *fnrl* mutant alleles (Fig. 2A,B). These results indicate that FNRL plays a role in ROS accumulation both under normal conditions and during isoxaben treatment.

**Figure 2:**
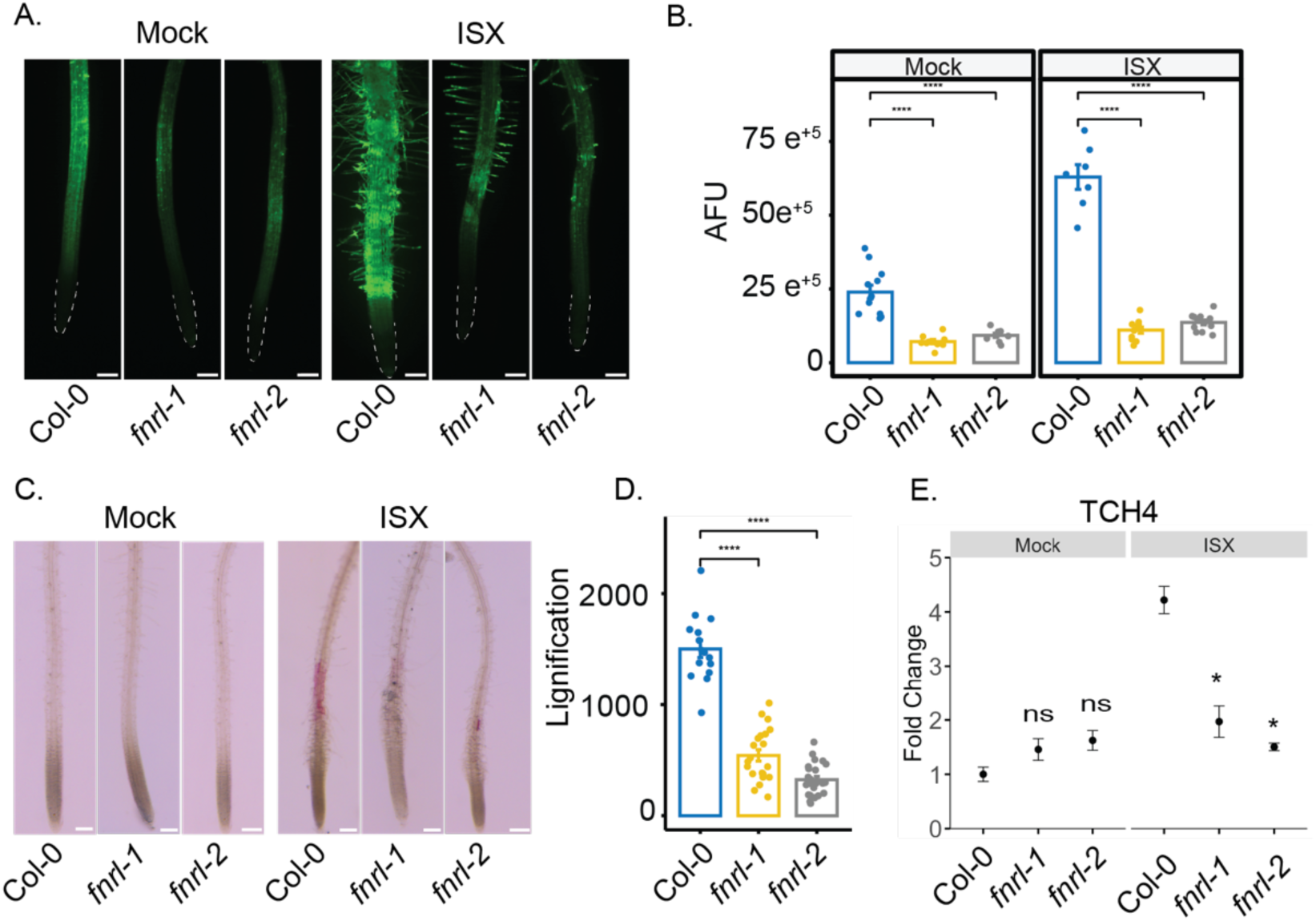
FNRL is involved in isoxaben-induced cell wall damage response. A) Representative images demonstrating ROS activity through H2DCFDA signal in the root tips of ten-day-old seedlings exposed to 2.5 nM isoxaben (ISX) or Mock conditions. Scale bar is 200 µm. B) Fluorescence quantification of the results presented in panel A in arbitrary fluorescence units (AFU). Quantification was performed in 2 mm sections of the root starting from the root tips (outlined). C) Phloroglucinol signal intensity indicating lignin accumulation in roots of ten-day-old seedlings exposed to 600 nM isoxaben (ISX) or Mock conditions for 6 h in freshly prepared medium. Scale bar is 200 µm. D) Quantification of the results presented in panel C. Quantification was performed in 2 mm sections of the root starting from the root tips. E) Gene expression levels of the isoxaben marker gene *TCH4* determined by qRT-PCR in ten-days-old seedlings exposed to 2.5 nM isoxaben (ISX) or Mock conditions. The results from three biological replicates are shown. Mean ± SD is presented. Statistical significance is denoted by asterisks (*, P < 0.05; **, P < 0.01; ***, P < 0.001; and ****, P < 0.0001) according to Student’s t-test. The graphics and statistical analysis were performed using the ggpubR package in R.

Isoxaben-induced CWD leads to alterations in cell wall biochemistry, characterised by ectopic lignification and callose deposition (Engelsdorf et al., 2018). We examined lignin accumulation in roots using phloroglucinol staining in ten-day-old seedlings grown on 1/2 MS liquid media. In comparison to the wild type, we observed reduced phloroglucinol staining in the root elongation zone of *fnrl* seedlings, indicating a decrease in ectopic lignification in response to isoxaben (Fig. 2C,D).

To investigate whether FNRL activity is required for the transcriptional regulation of specific marker genes in response to isoxaben-induced CWD, we next performed quantitative real-time polymerase chain reaction (qRT-PCR) experiments using RNA isolated from ten-day-old roots. We focused on TOUCH4 (TCH4), which is a marker gene that responds to isoxaben-induced CWD (Van der Does et al., 2017). We found that the isoxaben-induced upregulation of TCH4 was notably attenuated in the *fnrl* mutants when compared to the wild type (Fig. 2E). Taken together, FNRL appears to promote CWD-responses when seedling roots are exposed to isoxaben.

### *fnrl* mutants do not restore root growth of cellulose deficient mutant *prc1-1*

We next aimed to investigate whether *fnrl* mutants have a more general role in response to cellulose deficiency. We utilised the *prc1-1* mutant, which exhibits reduced cellulose due to a loss-of-function mutation in CESA6/PRC1 (Fagard et al., 2000), which causes impaired root elongation. To assess the influence of FNRL on *prc1-1*, we generated *prc1-1 fnrl-2* double mutants and compared their root length with that of *fnrl-2* and *prc1-1* single mutants (Fig. 3A). Intriguingly, the *prc1-1 fnrl-2* double mutant displayed a markedly shorter root length compared to the *prc1-1* single mutant (Fig. 3A,B), demonstrating an additive phenotype. These results indicate that FNRL and CESA6/PRC1 independently regulate root growth, and FNRL is unlikely to be involved in the cellulose-related aspects of isoxaben phytotoxicity during root growth.

**Figure 3:**
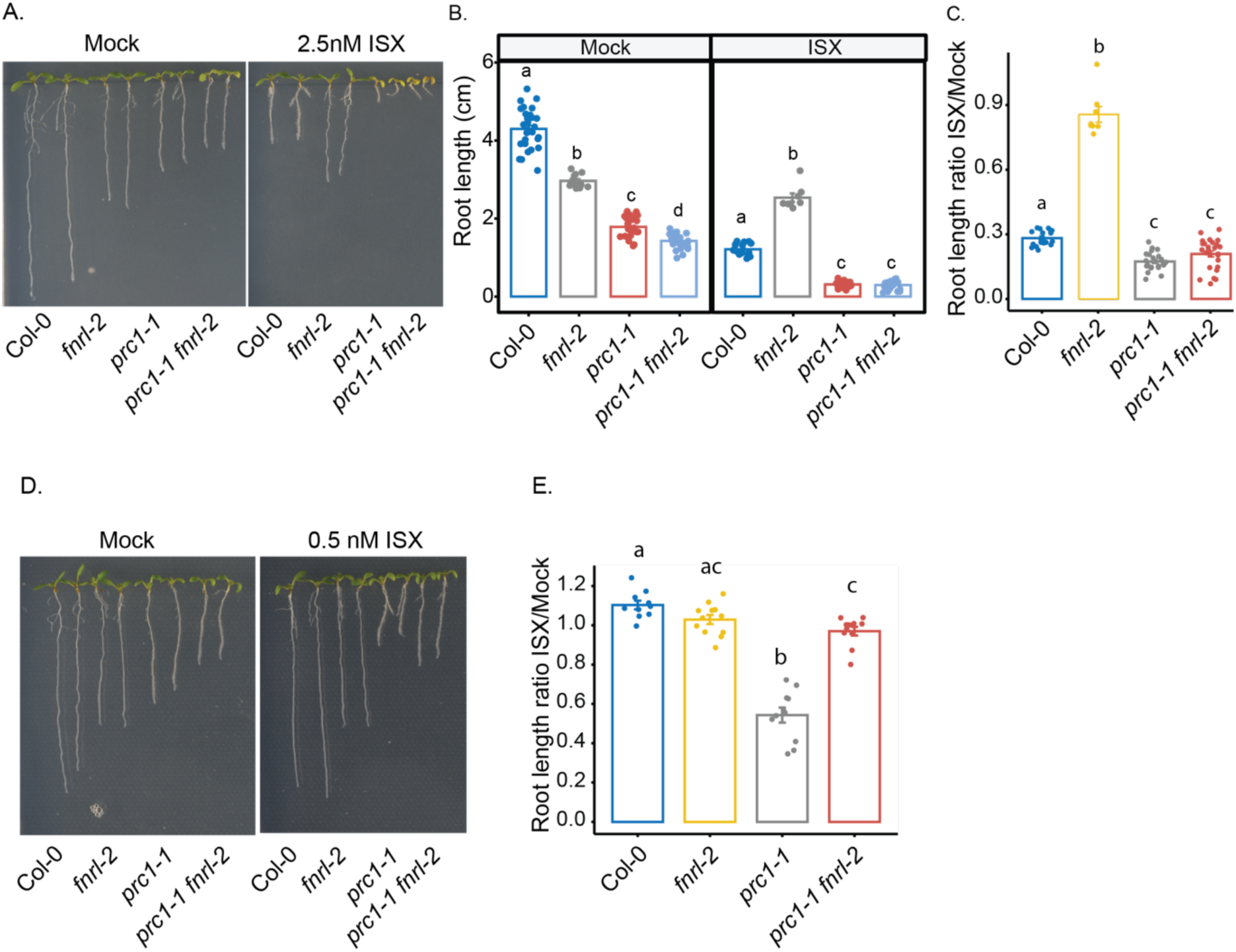
*fnrl* mutants fail to restore root growth in cellulose-deficient mutant *prc1-1*. A) Representative images of ten-day-old seedlings grown on 1/2 MS medium supplemented with 1% sucrose and 0.8% agar on Mock or 2.5 nM isoxaben (ISX) treatment. B) Quantification of primary root length in ten-day-old seedlings grown on 1/2 MS medium supplemented with 1% sucrose and 0.8% agar on Mock or 2.5 nM isoxaben (ISX) treatment. C) The ratio of primary root length reduction in response to isoxaben from the data presented in A. Ratio is calculated by averaging the Mock and dividing each isoxaben value by the Mock average. D) Representative images of ten-day-old seedlings grown on 1/2 MS medium supplemented with 1% sucrose and 0.8% agar on Mock or 0.5 nM isoxaben (ISX) treatment. E) The ratio of primary root length reduction in response to isoxaben from the data presented in D. Ratio is calculated by averaging the Mock and dividing each isoxaben value by the Mock average. For B, C and E, assigned letters (“a”, “b”, and “c”) signify distinct groups with significant differences between genotypes, calculated with one-way ANOVA followed by Tukey HSD test.

We next treated the mutant combinations with 2.5 nM isoxaben. The *prc1-1* mutant exhibited heightened sensitivity to isoxaben, a trait similarly observed in the *prc1-1 fnrl-2* double mutants (Fig. 3A,B,C). However, these plants deteriorated substantially, potentially obscuring any differences between the *prc1-1* and *prc1-1 fnrl-2* double mutants. To address this, we employed a lower dosage of isoxaben (0.5 nM). While *prc1-1* mutants exhibited increased sensitivity even at the lower isoxaben dose, the *prc1-1 fnrl-2* double mutants remained unresponsive to isoxaben (Fig. 3D,E). These findings illustrate the capacity of *fnrl* mutants to tolerate isoxaben, even in the genetically sensitive background of *prc1-1*.

### Isoxaben-induced transcriptomic changes are attenuated in *fnrl* mutants

To investigate whether other aspects of isoxaben-induced changes are evident in *fnrl* mutants, we performed RNA sequencing analyses on root tips 48 hours after isoxaben treatment. Ten-day-old plants were grown on 1/2 MS medium and then transferred to plates containing 2.5 nM isoxaben or mock for 48 hours. RNA was extracted from 2 mm sections of root tips, followed by RNA sequencing, which revealed distinct gene regulation patterns in response to isoxaben treatment between wild-type and *fnrl* seedlings.

Principal component analysis (PCA) plots demonstrated a clear separation of wild-type seedlings treated with isoxaben compared to mock (Fig. 4A). By contrast, *fnrl* mutants clustered together regardless of the treatment (Fig. 4A). Differentially expressed gene (DEG) analysis showed 444 differentially regulated genes in wild-type plants upon isoxaben treatment, whereas *fnrl-1* and *fnrl-2* mutants showed only five and four genes, respectively (Fig. 4B). These results suggest that *fnrl* mutants do not respond to isoxaben at the transcriptomic level. Gene ontology (GO) term enrichment analysis provided insights into the impact of isoxaben treatment on gene expression in wild type plants. Upregulated genes in wild type were associated with JA signalling, response to toxic substances, response to darkness, and root hair development, while downregulated genes were related to photosynthesis, photosystem, cyclin-dependent kinases, and secondary cell walls (Fig. 4C).

**Figure 4:**
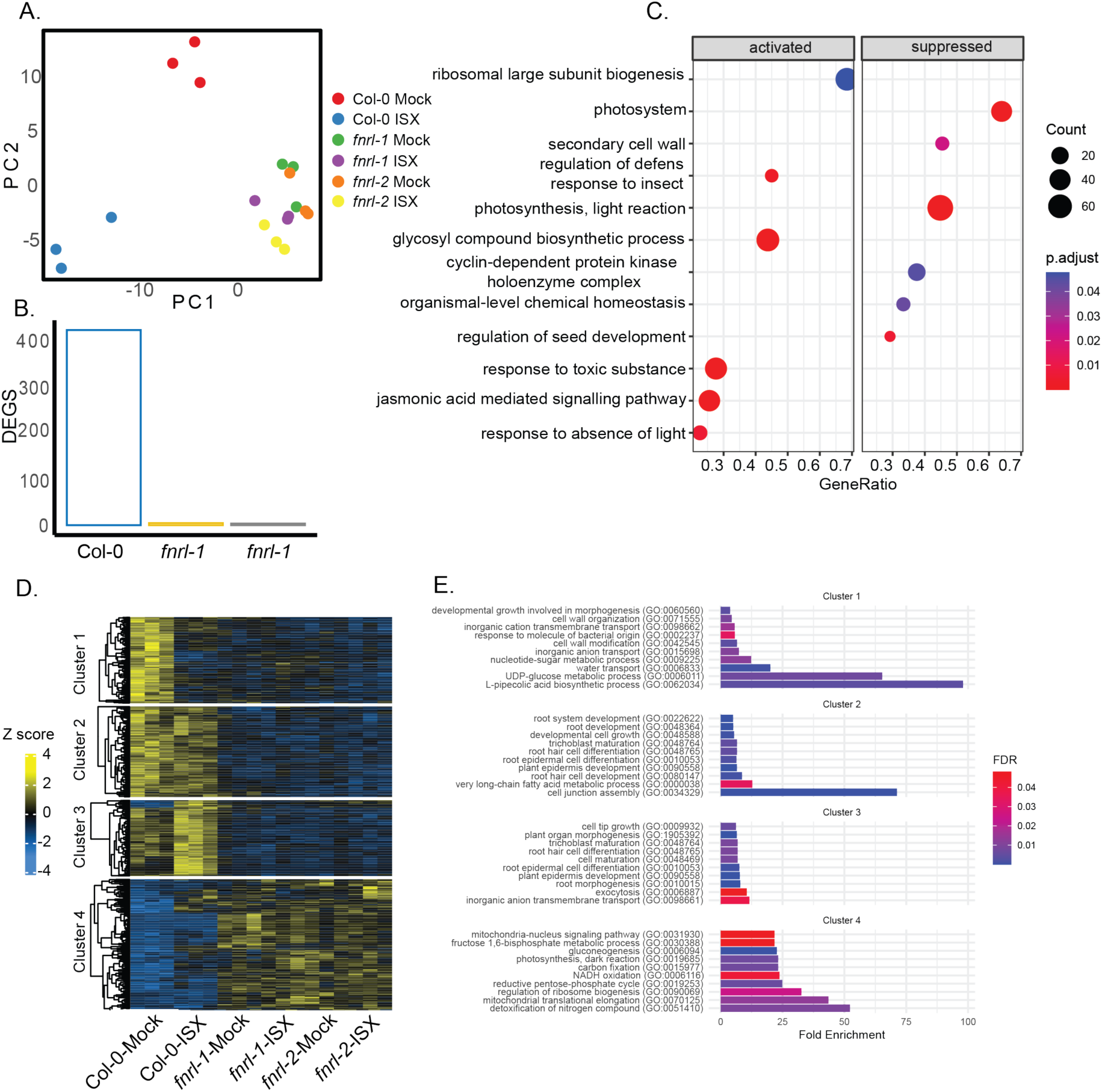
Whole transcriptomic profiling reveals impaired response of *fnrl* mutants to isoxaben treatment. A) PCA plot showing the separation of wild-type plants treated with isoxaben (ISX) from mock-treated plants. RNA was extracted from root tips after 48 h of exposure to 2.5 nm isoxaben or Mock treatment. B) Number of DEGs identified in response to isoxaben treatment in wild-type plants and *fnrl-1* and *fnrl-2* mutants using stringent criteria (log2FoldChange >1 or <-1 and padj<0.05). C) Bubble plot illustrating GO term enrichment analysis for DEGs in wild-type plants treated with isoxaben compared to mock-treated plants. D) Heatmap depicting hierarchical clustering of DEGs using the Ward method, resulting in the identification of four clusters. E) GO term enrichment analysis for the four identified DEG clusters in D.

Clustering analysis of DEGs revealed distinct isoxaben-response patterns (Fig. 4D). Cluster 1 consisted of genes that were downregulated by isoxaben and that were already downregulated in *fnrl* mutants under mock conditions. These genes were associated with GO terms related to UDP-glucose biosynthesis, cell wall organisation, and anion transport. This indicates that *fnrl* mutants may already exhibit isoxaben-like stress under mock conditions. Cluster 2 comprised genes that did not change with isoxaben treatment but were already downregulated in *fnrl* mutants under mock and associated with GO terms such as fatty acid metabolism, cell junction assembly, and root development. Cluster 3 contained genes upregulated by isoxaben in wild type, but that were downregulated in *fnrl* mutants. These genes were associated with GO terms related to anion transport, root hair development, and exocytosis. Cluster 4 consisted of genes that did not change upon isoxaben exposure, but were upregulated in *fnrl* mutants, and were associated with GO terms such as detoxification of nitrogen compounds, mitochondria-nucleus signalling pathway, and NADH oxidation (Fig. 4D). These findings indicate that *fnrl* mutants already exhibited a transcriptomic change resembling that of isoxaben-stressed plants, suggesting that *fnrl* mutants are likely primed for isoxaben stress response.

### *fnrl* mutants exhibit constitutively active mitochondria-nucleus retrograde signalling response

The analysis of RNA sequencing data revealed the enrichment of GO terms related to mitochondria-nucleus retrograde signalling in genes constitutively upregulated in *fnrl* mutants (Fig. 4D). Notably, *fnrl-1* and *fnrl-*2 exhibited constitutive upregulation of 14 mitochondria-nucleus retrograde signalling marker genes (Fig. 5A). To validate these findings, we performed qRT-PCR and confirmed the upregulation of mitochondria-nucleus signalling marker genes, including *ANAC013*, *alternative oxidase 1A* (*AOX1a*), and *up-regulated by oxidative stress* (*UPOX*) (Fig. 5B), in *fnrl-1* and *fnrl-2* mutants under both mock and isoxaben treatment conditions. These results suggest that mitochondria-nucleus signalling is constitutively active in the *fnrl* mutants.

**Figure 5:**
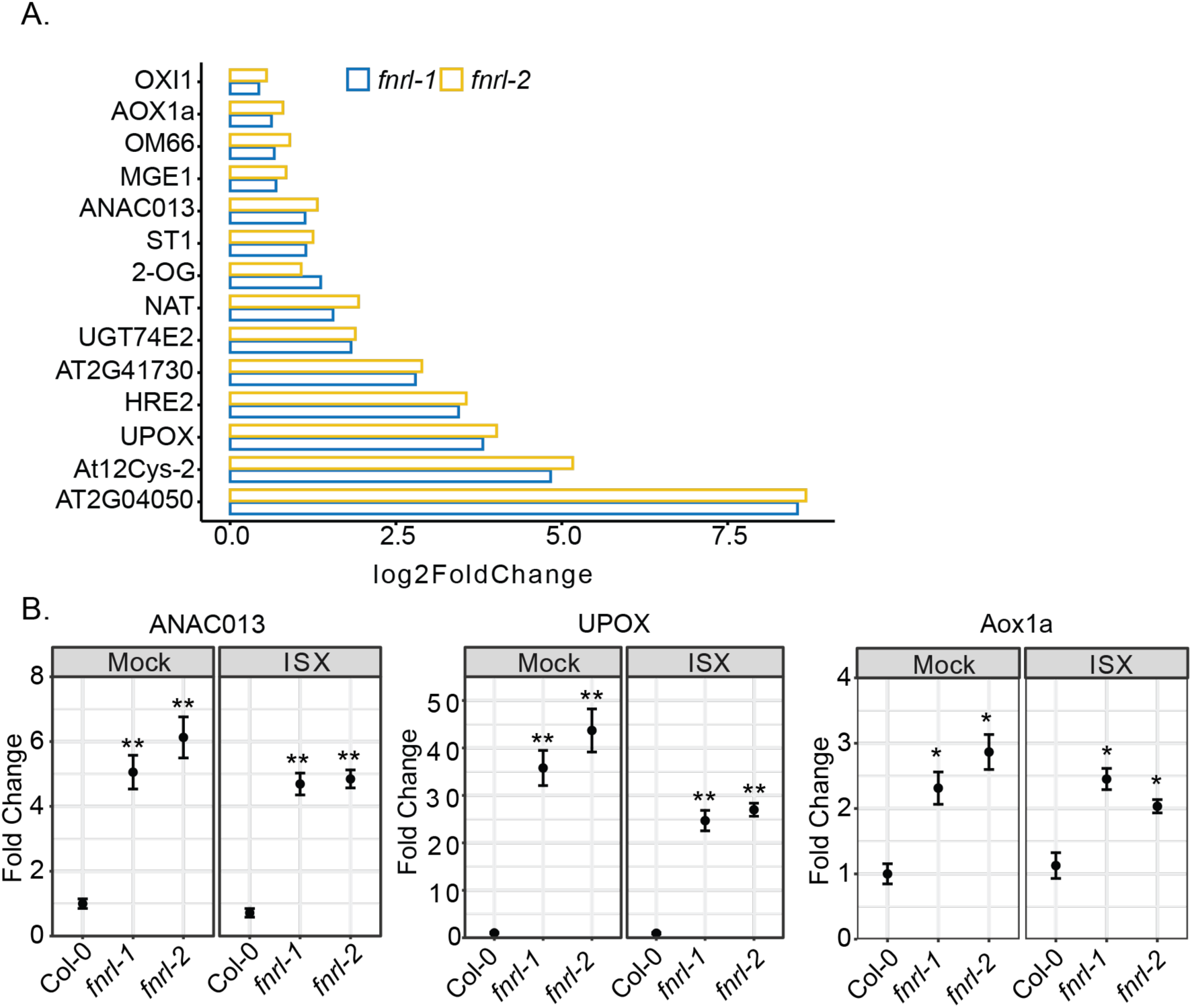
Activation of mitochondrial retrograde signalling response in *fnrl* mutants. A) RNA sequencing expression levels of genes belonging to the mitochondrial dysfunction stimulon, which share a common cis-element for the transcriptional response to mitochondrial stress, in *fnrl* mutant lines relative to wild type under mock treatment. Gene expression is presented as log2foldchange. B) Gene expression levels of mitochondria-nucleus retrograde signalling marker genes measured by qRT-PCR in ten-day-old seedlings treated with 2.5 nM isoxaben (ISX) or mock. The results are shown for three biological replicates. Marker genes and genotypes are indicated. Mean ± SD is presented. Statistical significance is indicated by asterisks (*, P < 0.05; **, P < 0.01; ***, P < 0.001; and ****, P < 0.0001) based on Student’s t-test.

FNRL is predicted to be plastid-localised (Koskela et al., 2018), so we also examined changes in plastid-nucleus signalling in the *fnrl* mutants. To do this, we performed a GENOME UNCOUPLED (GUN) assay (Zhao et al., 2018), where the mutants were grown on lincomycin and expression changes of multiple GUN phenotype marker genes were evaluated. The results indicated that *LIGHT-HARVESTING CHLOROPHYLL B-BINDING PROTEIN1.1* (*LHCB1.1)* and *CHLOROPLAST PROTEIN 12* (*CP12)* exhibited significantly higher expression in the control *gun1* mutants compared to the wild type when grown on 550 µM lincomycin, as expected (Zhao et al., 2018). However, the *fnrl* mutants displayed a response similar to the wild type (Supplemental Figure 2A,B,C), indicating that the GUN plastid retrograde signalling pathways are not active in *fnrl* mutants.

### ANAC017 plays a key role in *fnrl*-mediated isoxaben tolerance

The ANAC017 transcription factor is the master regulator of mitochondria-nucleus retrograde signalling (Ng et al., 2013). To investigate the involvement of ANAC017-dependent mitochondria-nucleus retrograde signalling in isoxaben tolerance, we generated *anac017-1 fnrl-2* double mutants, as well as *aox1a-1 fnrl-2* double mutants, which encodes an important downstream component of ANAC017 signalling, associated with mitochondrial dysfunction (Ng et al., 2013). The primary root length ratio of plants grown on isoxaben compared to the mock treatment revealed a pronounced tolerance in both alleles of *fnrl* mutants. Additionally, the *aox1a-1* mutant exhibited a subtle yet statistically significant tolerance to isoxaben, whereas the *anac017-1* mutant did not exhibit a significant deviation from the wild-type response. Interestingly, the loss of AOX1a did not affect the *fnrl*-mediated tolerance to isoxaben, as the roots of *aox1a-1 fnrl-2* double mutants remained tolerant to isoxaben treatment (Fig. 6A,B). However, compromising ANAC017 activity in the *anac017-1 fnrl-2* double mutant partially reduced isoxaben tolerance. This was evident through the shorter roots observed in the *anac017-1 fnrl-2* double mutant compared to the *fnrl-2* mutant in response to isoxaben treatment (Fig. 6A,B). These findings indicate that the activated mitochondria-nucleus signalling response in the *fnrl* mutants is partially responsible for isoxaben tolerance, and that ANAC017 plays a role in mediating the isoxaben tolerance conferred by *fnrl* mutants.

**Figure 6:**
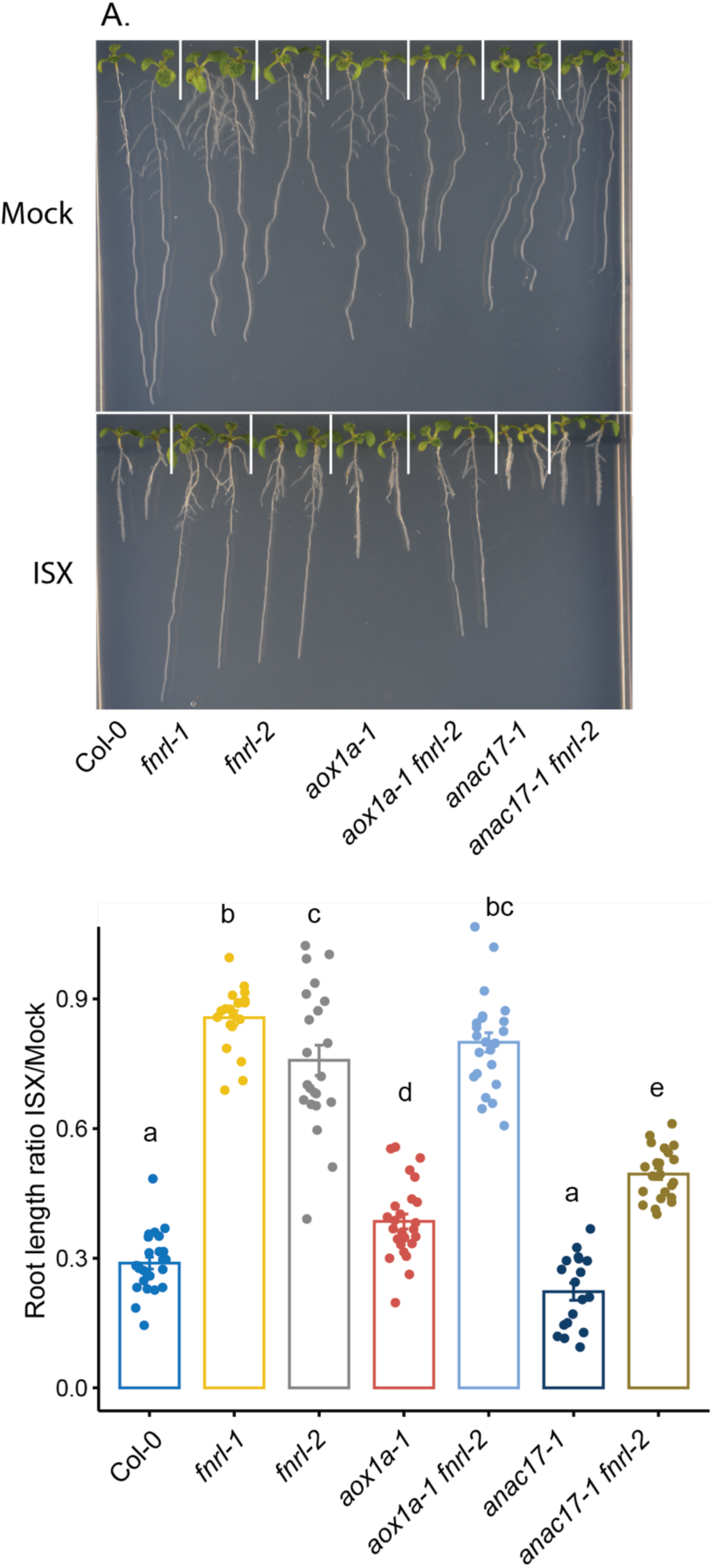
ANAC017 transcription factor plays a key role in *fnrl*-mediated isoxaben tolerance. A) Representative images of ten-day-old seedlings grown on 1/2 MS medium supplemented with 1% sucrose, 0.8% agar, and either 2.5 nM isoxaben (ISX) or Mock treatment. B) The ratio of primary root length in ten-day-old seedlings under 2.5 nM isoxaben or Mock conditions was calculated by averaging the Mock and dividing each isoxaben value by the Mock average. Assigned letters (“a”, “b”, and “c”) signify distinct groups with significant differences between genotypes, calculated with one-way ANOVA followed by Tukey HSD test.

### FNRL is highly conserved across the plant kingdom

Phylogenetic analysis of the FNRL protein across major viridiplantae lineages reveals its high conservation, typically existing as a single copy gene in dicotyledonous plants except for recent gene duplication events in species like *Populus trichocarpa* (Tuskan et al., 2006) and *Malus domestica* (Velasco et al., 2010). In dicotyledonous plants, the sequence similarity exceeds 80%, while monocotyledonous plants exhibit at least one homologue with a remarkably high similarity approaching 80%. A distinct clade is observed in monocotyledonous plants, with similarity ranging between 60 and 70%. The *FNRL* gene maintains high conservation in mosses and liverworts, with a similarity above 70%. Chlorophytes, however, exhibit the most distant homologues of *FNRL*, with a similarity between 55 and 62% (Supplemental Table 2).

Comparative alignment of canonical *FNR* genes in Arabidopsis, including the *Leaf FNR* (*LFNR1* and *LFNR2*) and *Root FNR* (*RFNR1* and *RFNR2*) genes, underscores their considerable divergence from all other sequences. This substantiates the distinctiveness of *FNRL* from the *FNR* genes. These findings show that the FNRL sequence is remarkably conserved throughout the Viridiplantae, suggesting functional conservation across diverse plant species.

The *prc1-1*, *anac017-1*, and *aox1a-1* mutants utilised in this study have been previously characterised (Fagard et al., 2000; Ng et al., 2013). The *fnrl* T-DNA insertion mutant lines, *fnrl-1* (SALK_039715) and *fnrl-2* (SALK_048627), were obtained from the Arabidopsis Biological Resource Center (ABRC, https://abrc.osu.edu/). To conduct the experiments involving isoxaben and mock treatments (equal volume of ethanol), a vertical agar plate method was employed. The standard media composition consisted of a half-strength MS Basal Salt mixture, 0.5% (w/v) MES (Sigma-Aldrich), 0.5% (w/v) sucrose (Sigma-Aldrich), and 0.8% (w/v) agar. For isoxaben treatment, the standard media was supplemented with 2.5 nM isoxaben (Sigma, 36138). Arabidopsis seeds were subjected to surface sterilisation using chlorine gas for three hours. Subsequently, they were resuspended in 0.1% (w/v) agarose, stratified at 4°C for two days, and then sown in a single row on 10-cm-square plates sealed with 3M Micropore tape. The plates were vertically arranged in racks within a growth chamber, which maintained a 16-hour light/8-hour dark cycle, a light intensity of 120 E m^-2^ s^-1^, and temperature conditions of 23 °C (day)/19 °C (night). The relative humidity inside the chamber was set to 60%.

## Discussion

In this study, we identified *FNRL* as the first plastid-localised gene involved in the response to the cell wall biosynthesis inhibiting herbicide, isoxaben. Loss-of-function *fnrl* mutants displayed strong tolerance to isoxaben, suppressing the typical root stunting response (Fig. 1). Furthermore, FNRL supressed lignin accumulation and the transcriptional regulation of isoxaben-responsive marker genes (Fig. 2B,E). These results indicate that *fnrl* mutants can supress the isoxaben response, similar to established genes involved in CWD response to CBIs or genetically induced cellulose deficiency. For example, THE1, a member of the CrRLK1L family, has been identified as a key regulator of CWI in response to cellulose deficiency (Hématy et al., 2007). THE1 plays a role in regulating stress-response genes, lignin, JA, and SA accumulation when seedlings are exposed to isoxaben (Engelsdorf et al., 2018). However, unlike THE1, which can alleviate the growth phenotype of the cellulose-deficient mutant *prc1-1* without restoring cellulose content (Hématy et al., 2007), *fnrl* mutants are unable to restore *prc1-1* growth (Fig. 3). Furthermore, *fnrl* mutants did not exhibit a reduction in cellulose upon exposure to isoxaben. These findings suggest that while THE1 functions downstream of cellulose deficiency in modulating the response to cellulose deficiency, FNRL appears to operate upstream of this process, inhibiting isoxaben-mediated cellulose reduction. In a similar manner, the leucine-rich repeat receptor kinase MIK2/LRR-KISS and receptor kinase STRUBBELIG (SUB) are also involved in mediating the response to isoxaben by regulating ROS and lignin accumulation. However, unlike *fnrl* mutants, *the1*, *mik2/lrr-kiss,* and *sub* mutants do not show tolerance to isoxaben-mediated growth inhibition (Van der Does et al., 2017; Chaudhary et al., 2020), suggesting that *fnrl*-mediated resistance to isoxaben displays unique features from previously known CWD response regulators.

Our RNA sequencing analysis showed that, in contrast to wild type, *fnrl* mutants were largely unresponsive to isoxaben. Comparing gene expression data between wild-type and *fnrl* mutants under mock conditions revealed significant differential regulation of numerous genes. Specifically, genes associated with anion transport were prominently downregulated in *fnrl* mutants (Fig. 4D). This downregulation of anion transporters could have profound implications for turgor, which is crucial in isoxaben-induced CWD response (Engelsdorf et al., 2018). Furthermore, we established that the mitochondria-nucleus retrograde signalling pathway is constitutively active in the *fnrl* mutants (Fig. 5). The FNRL protein is predicted to function as an NADPH dehydrogenase (Koskela et al., 2018), and likely plays a critical role in maintaining the balance of NADPH within chloroplasts. It may be anticipated that *fnrl* mutants would experience an accumulation of reducing equivalents in plastids. Consequently, the activation of the mitochondria-nucleus retrograde signalling pathway in *fnrl* mutants is likely a direct result of the putatively increased levels of reducing equivalents in plastids. To maintain cellular redox balance, cells employ the malate/oxaloacetate shuttle to transport excess reducing equivalents from plastids to the mitochondria (Taniguchi and Miyake, 2012; He et al., 2023). Subsequently, this increased influx of reducing equivalents in mitochondria activates the mitochondrial retrograde signalling pathway (Taniguchi and Miyake, 2012; He et al., 2023). While the interconnectedness of these organelles in cellular metabolism and stress response has been widely acknowledged (He et al., 2023), our characterisation of the *fnrl* mutants suggests that even seemingly mild plastid perturbation can activate the mitochondria-nucleus retrograde signalling pathway in root tissue. This highlights the intricacy of cellular communication and the ability of the cell to respond to perturbations in redox homeostasis through cross-organelle signalling mechanisms.

Depletion of transcription factor ANAC017, a master regulator of mitochondria-nucleus retrograde signalling, results in a reduction of the isoxaben tolerance phenotype triggered by *fnrl-2* (Fig. 6). This result is consistent with previous findings showing that ANAC017 regulates tolerance to another CBI, C17 (Hu et al., 2016). ANAC017 plays a crucial role in H2O2-mediated primary stress responses, particularly in retrograde signalling, accounting for more than 85% of such responses in plants (Ng et al., 2013). However, depleting the ANAC017 downstream target *AOX1a*, which plays a role in dissipating excess reducing equivalents in mitochondria, does not restore sensitivity of *fnrl* mutants to isoxaben (Fig. 6). This shows the role of ANAC017 in isoxaben tolerance is independent of AOX1a.

The expression of several uridine diphosphate (UDP) glycosyltransferases (UGTs) is significantly induced in *fnrl* mutants (Supplemental Table S1). UGTs are involved in catalysing the transfer of glycosyl groups to various lipophilic chemicals, neutralising their effects (Bock, 2016; Karlova et al., 2022). It is possible that isoxaben itself is glycosylated in plants overexpressing these UGTs, which could potentially contribute to isoxaben tolerance. This hypothesis would align with the RNA sequencing results where gene expression in *fnrl* mutants is almost unchanged after 48 hours of isoxaben exposure. Further investigation into isoxaben tolerance in UGT overexpressing lines and direct evidence of isoxaben glycosylation by these UGTs could shed further light on the underlying mechanisms of isoxaben tolerance.

Isoxaben stands out as a rare instance where no resistant weed species have emerged against its action in the field. For instance, wild radish (*Raphanus raphanistrum*) populations are becoming increasingly resistant to several commonly used herbicides from different modes of action (Ashworth et al., 2014; Owen et al., 2015) while pre-emergent applications of isoxaben can effectively control this weed. Despite its success in horticultural crops, isoxaben application is limited in agricultural crops.

Various isoxaben-tolerant *CESA* mutants and mutants in mitochondria localised proteins have been identified; however, the mutants typically display shoot growth penalties (Scheible et al., 2001; Desprez et al., 2002; Nakagawa and Sakurai, 2006; Shim et al., 2018), limiting their utility for breeding isoxaben tolerance traits. FNRL represents the first case of a plastid-localized protein whose mutants exhibit enhanced isoxaben tolerance. Moreover, the mechanism of isoxaben tolerance exhibited in the *fnrl* mutants stand out by lacking a noticeable shoot growth penalty (Fig. 1). Additionally, the high conservation of FNRL across all plant lineages (Fig. S3) implies the preservation of its function. Therefore, FNRL-mediated isoxaben tolerance has unique potential for biotechnological applications, particularly in developing isoxaben tolerance in crop plants. This can be achieved by targeting the FNRL gene using genome editing tools like CRISPR-Cas9, thereby expanding the herbicide’s utility.

## Methods

### Plant material and growth conditions

The *prc1-1*, *anac017-1*, and *aox1a-1* mutants utilised in this study have been previously characterised (Fagard et al., 2000; Ng et al., 2013). The *fnrl* T-DNA insertion mutant lines, *fnrl-1* (SALK_039715) and *fnrl-2* (SALK_048627), were obtained from the Arabidopsis Biological Resource Center (ABRC, https://abrc.osu.edu/). To conduct the experiments involving isoxaben and mock treatments (equal volume of ethanol), a vertical agar plate method was employed. The standard media composition consisted of a 1/2 MS Basal Salt mixture, 0.5% (w/v) MES (Sigma-Aldrich), 0.5% (w/v) sucrose (Sigma-Aldrich), and 0.8% (w/v) agar. For isoxaben treatment, the standard media was supplemented with 2.5 nM isoxaben (Sigma-Aldrich, 36138). Arabidopsis seeds were subjected to surface sterilisation using chlorine gas for three hours. Subsequently, they were resuspended in 0.1% (w/v) agarose, stratified at 4°C for two days, and then sown in a single row on 10-cm-square plates sealed with 3M Micropore tape. The plates were vertically arranged in racks within a growth chamber, which maintained a 16-hour light/8-hour dark cycle, a light intensity of 120 E m^-2^ s^-1^, and temperature conditions of 23 °C (day)/19 °C (night). The relative humidity inside the chamber was set to 60%.

### qRT-PCR

qRT-PCR was performed as previously described (Khan et al., 2020). Briefly, upon harvesting, the roots were rapidly frozen in liquid nitrogen and subsequently ground using an Invitrogen TissueLyser. Approximately 100 mg of the resulting powder was employed to extract total RNA, utilising an RNeasy Plant Mini Kit (QIAGEN). To eliminate any potential DNA contamination, on-column DNase treatment was conducted using an RNase-free DNase kit (QIAGEN). For reverse transcription, 500 ng of RNA per sample was utilised in conjunction with SuperScript III reverse transcriptase from Invitrogen. Subsequently, qRT-PCR was performed using the SYBR select master mix from Invitrogen according to the manufacturer’s instructions. The primers employed for the qRT-PCR analysis are described in Supplemental Table 3.

### ROS and lignin staining and cellulose quantification

To assess intracellular ROS accumulation in root meristems, H2DCFDA fluorescent stain (Sigma-Aldrich, D6883) was employed through a modified version of an established protocol (Juárez et al., 2015). Arabidopsis seeds were cultivated on square plates containing 1/2 MS medium supplemented with 1% sucrose, 0.8% agar, and 2.5 nM isoxaben or mock treatment. After stratification at 4°C for two days, the seedlings were grown for ten days at 22°C under long-day conditions with a 10-degree incline. For imaging, the seedlings were treated with 100 μM H2DCFDA staining solution for 5 minutes and then rinsed with water on the same plates. Images were taken on a Leica M205 FCA stereo fluorescence microscope with a GFP ET filter. Lignin staining using phloroglucinol was conducted following a previously established protocol (Gigli-Bisceglia et al., 2018). Images of phloroglucinol-stained seedlings were taken on a Leica M205 FCA stereo fluorescence microscope. Micrographs of ROS and lignin staining were quantified using ImageJ software (Rueden et al., 2017). For quantification, a defined region of interest positioned 500 μm above the root tip (excluding the root cap) was selected for all samples. Cellulose quantification was performed as previously described (Khan et al., 2023).

### RNA sequencing and data analysis

Seedlings were grown for eight days on 1/2 MS media and then transferred to mock- or 2.5 nM isoxaben-containing plates. Two mm root tips were harvested after 48 hours of treatment. Three biological replicates were used for each genotype and treatment. The replicates consisted of 2 mm root tip sections obtained from more than one hundred pooled seedlings from at least two independent plates. Total RNA was extracted from these replicates using an RNeasy Plant Mini Kit. The TruSeq Stranded mRNA Library Prep Kit was employed to generate RNA-seq libraries, which were subsequently sequenced on an in-house Illumina NextSeq500 instrument. The resulting reads were 70 base pairs in length. To ensure the quality and reliability of the data, FastQC software (https://www.bioinformatics.babraham.ac.uk/projects/fastqc/) was used for quality control assessment of the data to verify its integrity. Prior to alignment, the RNA-Seq reads were trimmed using Trimmomatic (Bolger et al., 2014). The trimmed reads were aligned to the Arabidopsis TAIR10 genome employing STAR (v2.5.3) (Dobin et al., 2013). For each gene, the htseq-count tool was used to count the mapped reads (Anders et al., 2015). The htseq count output served as the input for DESeq2 analysis to identify differentially expressed genes (Love et al., 2014). Gene clustering was performed with the clusterprofiler package in R (Wu et al., 2021). Heatmaps were drawn using a complex-heatmap package (Gu, 2022) and the GO enrichment was performed on geneontology.org. The raw data have been deposited in the sequence read archive (SRA) under the BioProject accession number PRJNA1054327.

### Phylogenetic analysis

Phylogenetic analyses were conducted by utilizing the FNRL amino acid sequence to search for closest homologues in Phytozome (https://phytozome-next.jgi.doe.gov/) from 62 different species spread across major lineages of Viridiplantae. The similarity percentage and score for these sequences were retrieved from Phytozome. The identified protein sequences (Supplementary Table 1) along with canonical FNRs from Arabidopsis including LFNR1, LFNR2, RFNR1, and RFNR2 underwent alignment followed by the construction of a phylogenetic tree using NGphylogeny (https://ngphylogeny.fr/) using one click method. The resultant tree was then imported into iTOL (https://itol.embl.de/upload.cgi), from which the unrooted tree was obtained. Major clades were coloured and slight aesthetic modifications were performed in Adobe Illustrator.

## Accession numbers

FNRL: AT1G15140, ANAC017: AT1G34190, AOX1a: AT3G22370, GUN1: AT2G31400, TCH4: AT5G57560, RFNR1: AT4G05390, RFNR2: AT1G30510, LFNR1: AT5G66190, LFNR2: AT1G20020, PRC1: AT5G64740

## Supplemental data

Supplemental Table 1. Full dataset of gene expression analysis in the root tips of wild type, *fnrl-1,* and *fnrl-2* mutants in response to 48 hours of 2.5 nM isoxaben or Mock treatment.

Supplemental Table 2. List of FNRL homologues identified in diverse plant lineages and their similarity scores.

Supplemental Table 3. Primers used in this study.

