## Supplemental figures for "The *fnr-like* mutants enhance isoxaben tolerance by initiating mitochondrial retrograde signalling"

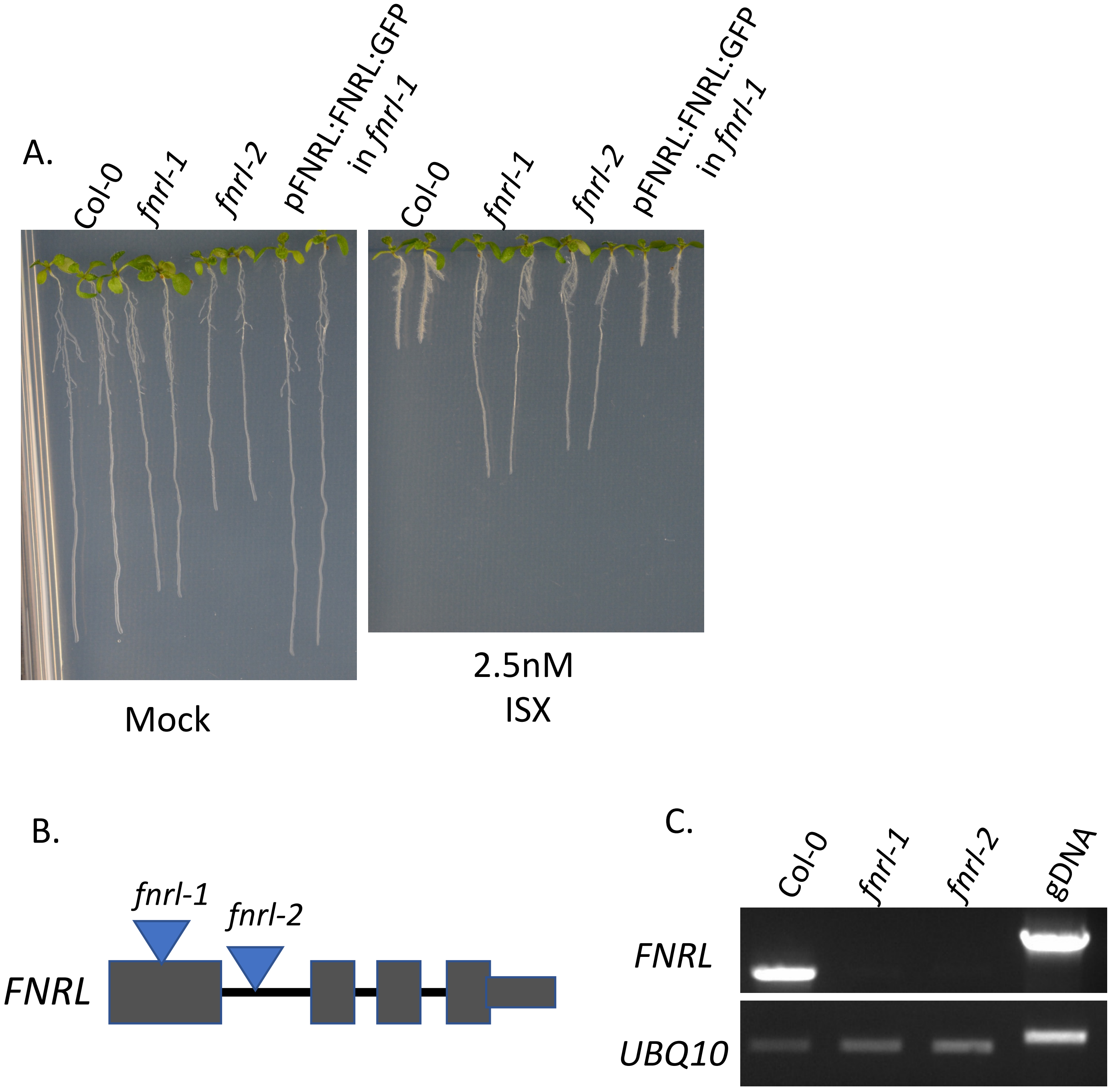


**Supplemental Figure 1: *fnrl-1* complementation, T-DNA insertion sites, and *FNRL* expression in *fnrl* mutants**

(A) Root length phenotype of indicated genotypes grown on Mock and 2.5 nM isoxaben (ISX) media for 10 days.

(B) T-DNA insertion sites of *fnrl-1* and *fnrl-2* alleles in the *FNRL* gene.

(C) Expression of *FNRL* in *fnrl-1* and *fnrl-2* mutants compared to the wild type. RNA was extracted from the roots of 7-day-old seedlings. Reverse transcription was performed on 1 µg of RNA, followed by standard PCR using full-length *FNRL* primers spanning from the ATG start codon to the stop codon. UBQ10 primers were used as a control, and genomic DNA (gDNA) was used as a control for gDNA contamination.

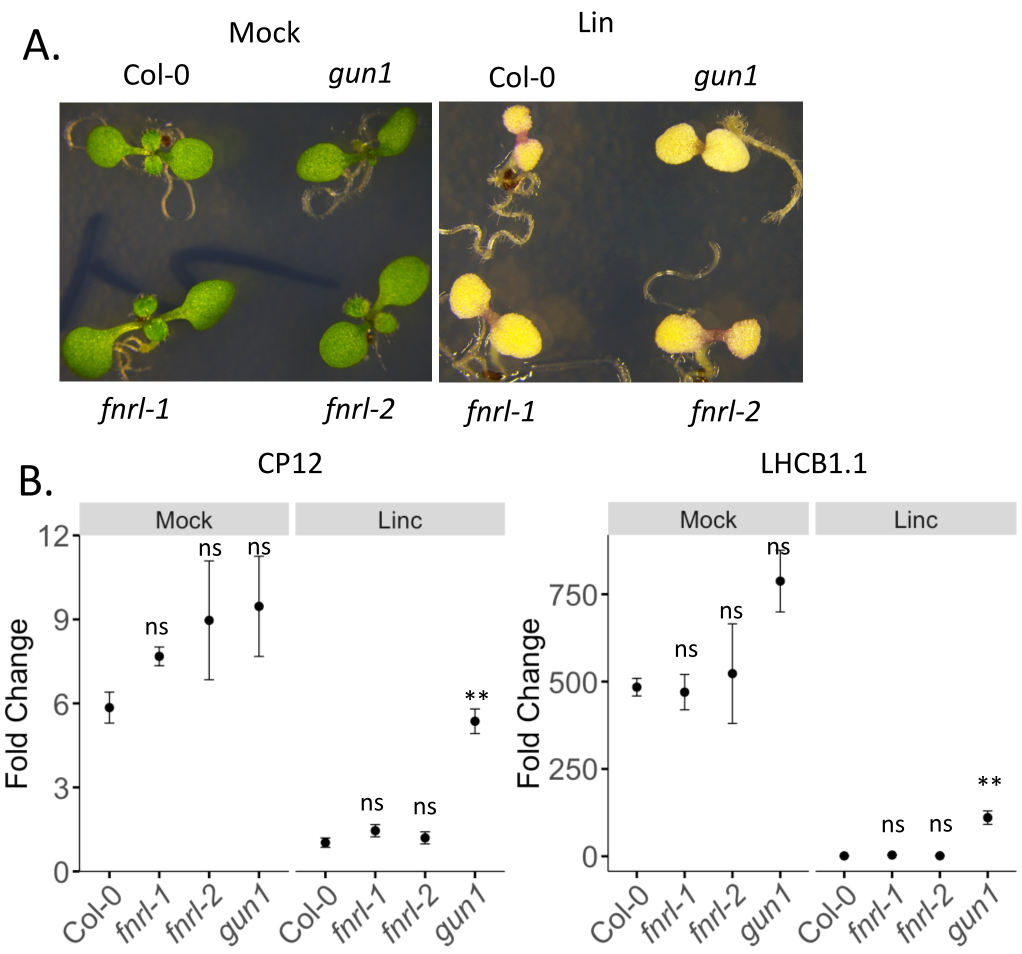


**Supplemental Figure 2: chloroplast retrograde signalling pathways are not active in the *fnrl* mutants.**

A) Representative images of Arabidopsis wild type (Col-0), *gun1*, *fnrl-1*, and *fnrl-2* seedlings grown in the absence or presence of 550 µM lincomycin (Linc).

B) Gene expression levels of chloroplast retrograde signalling marker genes measured by qRT-PCR in seven-day-old seedlings treated with 550 µM lincomycin or mock. The results are shown for three biological replicates. Mean ± SD is presented. Statistical significance is indicated by asterisks (ns, non-significant; *, $P < 0.05$; and **, $P < 0.01$) based on Student's t-test.

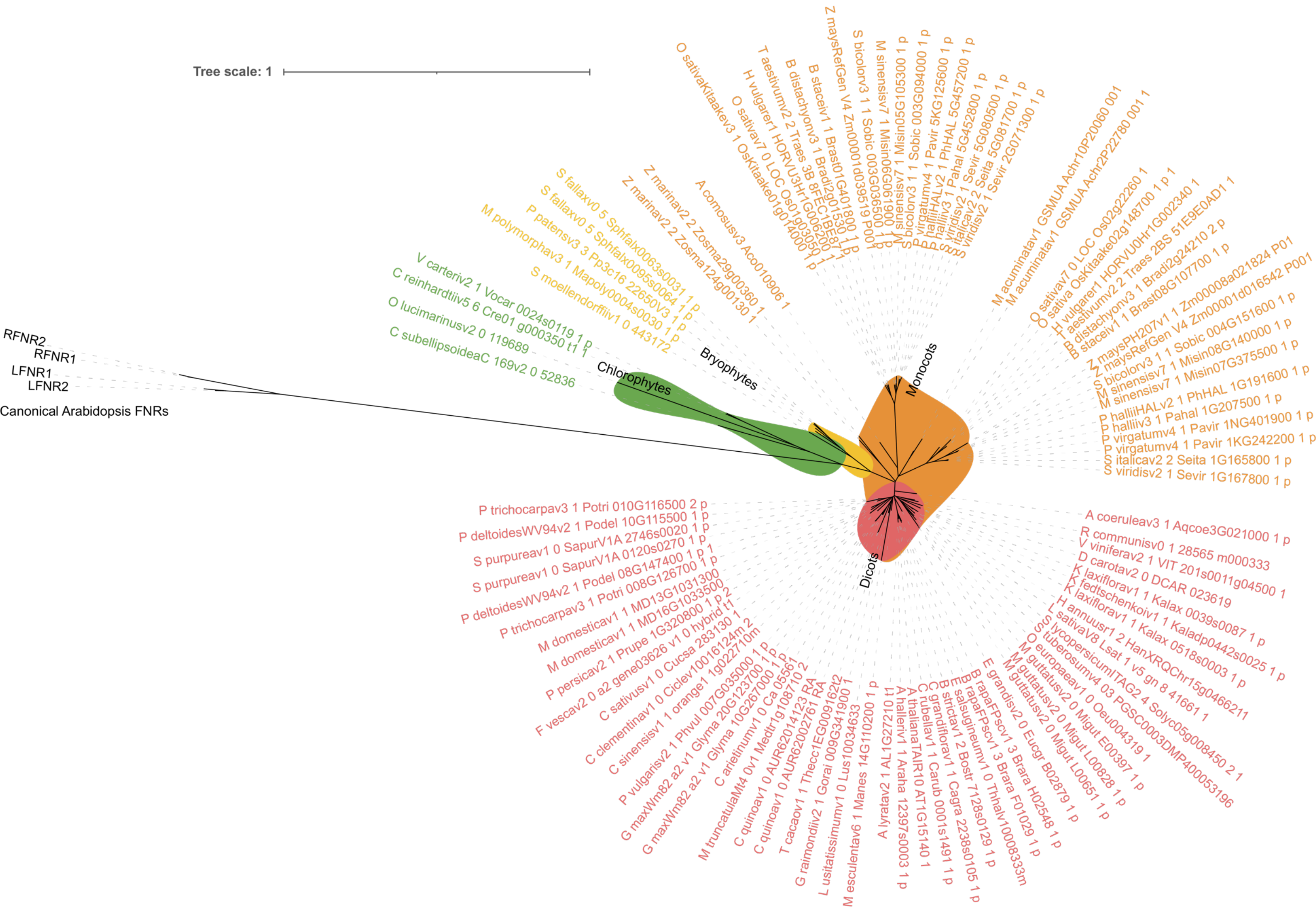


**Supplemental Figure 3: Phylogenetic analysis of FNRL homologues in plant lineages.**
Unrooted phylogenetic tree of FNRL homologues and FNR genes across diverse plant lineages. We conducted phylogenetic analyses using the FNRL amino acid sequence, searching for homologues in Phytozome across 62 Viridiplantae species. Sequences, including canonical FNRs from Arabidopsis, underwent alignment and tree construction using NGphylogeny. The resulting tree was visualized in iTOL and minor aesthetic adjustments were made in Adobe Illustrator.
